# Correlation network analysis based on untargeted LC-MS profiles of cocoa reveals processing stage and origin country

**DOI:** 10.1101/2020.02.09.940585

**Authors:** Santhust Kumar, Roy N. D’Souza, Marcello Corno, Matthias S. Ullrich, Nikolai Kuhnert, Marc-Thorsten Hütt

## Abstract

In order to implement quality control measures and create fine flavor products, an important objective in cocoa processing industry is to realize standards for characterization of cocoa raw materials, intermediate and finished products with respect to their processing stages and countries of origin. Towards this end, various works have studied separability or distinguishability of cocoa samples belonging to various processing stages in a typical cocoa processing pipeline or to different origins. Limited amount of success has been possible in this direction in that unfermented and fermented cocoa samples have been shown to group into separate clusters in PCA. However, a clear clustering with respect to the country of origin has remained elusive. In this work we suggest an alternative approach to this problem through the framework of correlation networks. For 140 cocoa samples belonging to eight countries and three progressive stages in a typical cocoa processing pipeline we compute pairwise Spearman and Pearson correlation coefficients based on the LC-MS profiles and derive correlation networks by retaining only correlations higher than a threshold. Progressively increasing this threshold reveals, first, processing stage (or sample type) modules (or network clusters) at low and intermediate values of correlation threshold and then country specific modules at high correlation thresholds. We present both qualitative and quantitative evidence through network visualization and node connectivity statistics. Besides demonstrating separability of the two data properties via this network-based method, our work suggests a new approach for studying classification of cocoa samples with nested attributes of processing stage sample types and country of origin along with possibility of including additional factors, e.g., hybrid variety, etc. in the analysis.

## 1. Introduction

Cocoa, scientifically *Theobroma cacao*, is a commodity of commercial interest to farmers as a crop and to businesses as a raw material for producing various cocoa based food products. Therefore, quality, variety and characteristics of cocoa and its derived food items have become an important area of research and development. Quality control (Fayeulle et al., 2019; Guehi et al., 2010; Kongor et al., 2016; Lima et al., 2011) and design of single origin cocoa products (Oberrauter et al., 2018; Ozretic-Dosen et al., 2007) are two of many focus areas in cocoa research. The former helps in ensuring whether the stages in a typical cocoa processing pipeline have been rightly carried to achieve the best possible finished product, and the latter commands high value among consumers for nuanced taste and aroma of the consumed food item.

Previous research successfully demonstrate characteristic differences between unfermented, partially fermented and fermented cocoa samples (processing-stages) and even identified corresponding potentially responsible classes of compounds through multivariate statistical analysis, e.g., principal component analysis (PCA) (Wold et al., 1987), on the chemical composition of these samples (Caligiani et al., 2014; D’Souza et al., 2017; Kumari et al., 2018; Megías-Pérez et al., 2018). Baring a few cases where the number of distinct countries relating to the samples in dataset at hand is few (D’Souza et al., 2017; Milev et al., 2014; Oliveira et al., 2016) or based on large continental regions (Acierno et al., 2016, 2018; Bertoldi et al., 2016; Kumari et al., 2018; Marseglia et al., 2016), a successful characteristic differentiation amongst samples on the basis of their country of origin has remained hard to define through metabolomic analysis (D’Souza et al., 2017; Sirbu et al., 2018; Vázquez-Ovando et al., 2015).

On the other hand, the ‘language of networks’ (Albert and Barabási, 2002; Newman, 2003) has proven immensely useful in visualizing and interpreting relationships between multitude of entities, and across many disciplines—metabolomics (Jeong et al., 2000), genetics (Grimbs et al., 2019; Kumar et al., 2018), proteomics (Szklarczyk et al., 2015), social science (Borgatti et al., 2009), logistics (Becker et al., 2012), gut ecology (Claussen et al., 2017) medicine (Barabási et al., 2011; Batushansky et al., 2016), finance (Kumar and Deo, 2012; Namaki et al., 2011), etc. to name a few. Some works have successfully applied this approach in the field of food science (Ahn et al., 2011; Hochberg et al., 2013; Ursem et al., 2008; Wang et al., 2017). Here, we apply the framework of network science to simultaneously study the clustering of cocoa samples with regards to their processing-stage sample types and country of origin.

We start by computing pairwise Spearman and Pearson correlation coefficients between 140 cocoa samples belonging to three different stages in a typical cocoa processing pipeline (unfermented, fermented and liquor) and 8 countries through their LC-MS profiles in positive ion mode. On the basis of correlations obtained, we construct correlation networks, at varying correlation thresholds. In these networks, the nodes are samples and an edge between two samples is drawn, when the correlation coefficient exceeds the threshold value.

We find that, as we progressively increase the correlation threshold from 0 towards 1, the clustering of cocoa samples is first dominated by processing-stage sample types at low and intermediate correlation thresholds, and then by countries of origin at high correlation thresholds. We show this both qualitatively and quantitatively via network visualizations and network edge statistics.

Our work demonstrates the presence of processing-stage level grouping on a coarser level and origin level grouping on a finer level within the former. This nested grouping can be revealed by successively keeping higher correlations. Further, our works suggests a new approach to study clustering or classification of food samples upon multiple nested attributes and can prove an important complement to traditional approaches and strategies.

## 2. Materials and Methods

### 2.1 Country and Origin details

The LC-MS data set we use here has a total of 140 samples (positive ion mode). The samples have been gathered and their LC-MS profiling done under COMETA project over a range of about past five years. These samples can be grouped into three sample-types (Unfermented, Fermented and Liquors) and eight origins (Brazil, Cameroon, Ecuador, Ghana, Indonesia, Ivory Coast, Malaysia and Tanzania). A cross-table of details about number of samples belonging to particular sample-type and country is given in Table 1.

**Table 1.** Sample division. The LCMS data set can be grouped on twin axes: sample-type and origin. There are 3 sample-types: Unfermented, Fermented and Liquors, and there are 8 origins (Brazil, Cameroon, Ecuador, Ghana, Indonesia, Ivory Coast, Malaysia and Tanzania).

|  | Brazil | Cameroon | Ecuador | Ghana | Indonesia | Ivory<br>coast | Malaysia | Tanzania | All |
| --- | --- | --- | --- | --- | --- | --- | --- | --- | --- |
| Unfermented | 4 | 3 | 8 | 0 | 14 | 16 | 6 | 3 | 54 |
| Fermented | 4 | 3 | 12 | 0 | 16 | 16 | 3 | 9 | 63 |
| Liquor | 0 | 6 | 3 | 5 | 0 | 9 | 0 | 0 | 23 |
| All | 8 | 12 | 23 | 5 | 30 | 41 | 9 | 12 | 140 |

### 2.2 Data pre-processing and cleaning

The data generation, cleaning, standardization and organization has been discussed in an earlier work (Kumar et al *previous manuscript*). Briefly, LC-MS data of all the samples was processed using MZMine (Pluskal et al., 2010) giving peak area list and corresponding *m/z ratio* and *retention times*. The detected compounds are assigned names/chemical formula on the basis of four ionization states ([M+H], [M+2H], [M+3H], [2M+H]) when possible, else the compound was named as ‘Unknown_’ suffixed with the *m/z* value, e.g., Unknown_865.1927. The samples were then put in an excel file, where each row represents a sample, and the column contain information about the sample-type, origin and peak areas of various compounds sorted in descending order by their mean peak are across all the samples.

### 2.3 Network production and visualization

Spearman and Pearson correlation analysis, and network generation/transformation was carried by writing programs from scratch in Python programming language making use of popular modules such as Pandas (McKinney, 2010, 2011) and NetworkX (Hagberg et al., 2008). Network visualization has been done in Cytoscape (Shannon et al., 2003). For layout of the network either of the following two variants of spring layout, which were available in Cytoscape itself, were used: (a) Edge-weighted Spring Embedded Layout (Kamada and Kawai, 1989), (b) Compound Spring Embedder (CoSE) (Dogrusoz et al., 2009). These layouts take into account the weight of the edge (in our case the Spearman or Pearson correlation coefficient) between nodes, so that the nodes with higher weight (correlations) are placed closer together.

### 2.4 Null model network or control network

A null model network is made by randomizing the weights (correlations) of edges in the original correlation network. It is important to note that the null model network so obtained has the same correlation distribution as that of the original correlation network because the set of correlations in the network remains unchanged, only the correlations between nodes is randomized. An ensemble of 100 such null model networks were generated. The reported statistics about a studied property on the null model networks is obtained by making calculations over this ensemble and then reporting the mean and standard deviation of the studied property. Higher the difference in the studied property between the original network and null network ensemble, higher the significance of the observed property in the original network.

## 3. Results

### 3.1 Correlation between cocoa samples

A typical LC-MS profile contains information about thousands of compounds present in a given sample defined by their retention time and associated *m/z* values (Kuhnert et al., 2013). Using the areas of peaks as a rough measure for concentration of these compounds across all samples, we calculate the Spearman and Pearson correlation coefficients (*r*) for all pairs of samples in our dataset.

The LC-MS data can be represented as a matrix L with entries 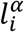. The upper index α represents the sample and lower index *i* represents the compound. Thus, the scalar quantity 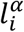 represents the concentration of *i*^th^ compound in the α^th^ LC-MS sample. Correspondingly, *l*^α^ is a vector which represents the LC-MS profile of sample α. The Pearson correlation between two LC-MS samples, say α and β with corresponding profiles *l*^α^ and *l*^β^, can be denoted as *r*_αβ_. It is calculated as

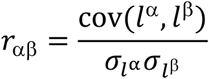

Where cov(*l*^α^, *l*^β^) represents the covariance between the LC-MS profiles of samples α and β, while 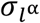 and 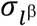 represent the standard deviation in the LC-MS profiles *l*^α^ and *l*^β^, respectively. The Spearman correlation can be defined as the Pearson correlation between the ranks of the original variables (i.e., *l*^α^ and *l*^β^). The ranked variables 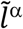 and 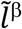, are obtained from the original variables *l*^α^ and *l*^β^ by sorting them from lowest to highest and substituting the values by the position in the sorted list (i.e., the rank of the values). Formally, the Spearman correlation coefficient is thus calculated as

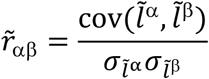

The Spearman and Pearson correlations across all pairs of LC-MS samples can be written in the form of matrices, 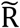 and R, whose entries denoted by 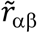 and *r*_αβ_, respectively.

The correlation matrices 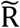 so obtained, i.e. the case of Spearman correlation coefficient, is visualized through heatmap in Figure 1A. The heat map of Pearson correlation coefficient matrix, R, is given in Supplementary Information file. By construction the correlation matrices 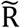 and R are symmetric. The twin attributes of nodes, namely the processing-stage sample type and country of origin, have been alternatively marked on the sides. Three blocks corresponding to Unfermented, Fermented and Liquor samples blocks are clearly distinguishable. It is also visible that Fermented and Liquor samples are part of a larger block which is separated from Unfermented samples. This shows that Liquor samples are closer in character to Fermented samples. This is in consonance with general expectation that liquor follows the fermentation stage. Furthermore, more chemical changes occur in cocoa when moving from unfermented stage to fermented stage than occurs from fermented to liquor stage. In case of correlation heatmap obtained using Pearson correlation (Supplementary Information file) the block of Unfermented samples is clearly distinguishable from Fermented and Liquor samples, while the Fermented and Liquor samples are mildly distinguishable. Further, it is important to note that no block structure on the basis of country is discernable at this level of detail about the correlations.

**Figure 1.**
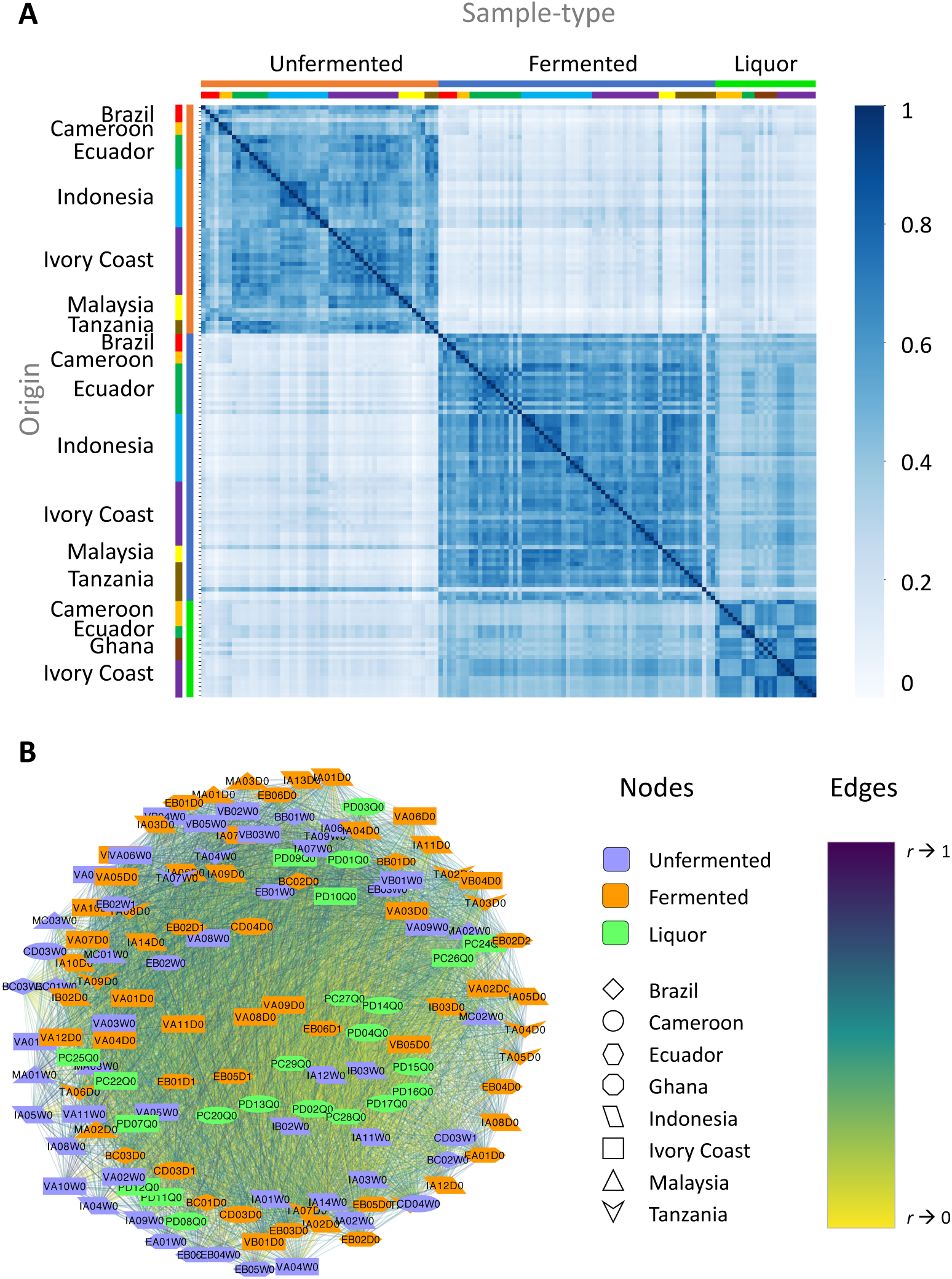
Correlation between cocoa samples. **(A)** Correlation heatmap. Darker regions represent high correlation, and lighter regions represent low correlation. Samples have been sorted on twin axes, first on processing stage sample-type, and then second internally on country of origin. Two distinct square block regions are clearly visible along the diagonal of the matrix, corresponding to Unfermented (smaller block) and Fermented (bigger block) samples. (**B)** Correlation Network. The correlation network made using all correlations between the set of cocoa samples using Spearman correlation. The nodes are color coded according to their processing-stage sample type and shape coded by their country of origin. The colors of edges code for the strength of correlation between nodes. The network is visualized using Cytoscape (Shannon et al., 2003) with ‘edge-weighted spring embedded layout’ which keeps nodes connected with higher correlations closer together.

Next, we define correlation network using the Spearman 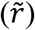 and Pearson correlations (*r*) obtained above. A network is defined through two sets of entities: nodes (*N*) and edges (*E*). The nodes denote the objects which are related to each other in some way, and the edge represent the relation between the nodes. For further knowledge about network, see (Albert and Barabási, 2002; Newman, 2003). In a correlation network, an edge represents the correlation between two nodes. In our correlation network, the nodes represent the different LC-MS samples of cocoa or its products sourced from different origins, and the edge between the nodes represent the correlation between the LC-MS samples. Figure 1B shows the correlation network obtained by using all correlations (0 to 1) between all LC-MS samples and visualized with edge-weighted spring layout (see section 2.3 Network production and visualization). Metadata about the LC-MS samples, such as country, and processing-stage sample type (unfermented, fermented, or liquor) has been represented through color and shape of nodes, respectively. The network shown in Figure 1B is the correlation network made using Spearman correlation and has 140 nodes and 6833 edges, i.e. 140 cocoa LC-MS samples and 6833 correlations 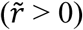 between the nodes. The network made using Pearson correlation is shown in the Supplementary Information file. The label of the node represents the internal LC-MS id. The strength of correlation is represented by the color of the edge between the nodes, yellow representing low correlation and violet representing high correlation. The spatial placement of nodes in Figure 1B, and all of the following networks, is done through variants of spring layout algorithms in Cytoscape (Shannon et al., 2003) which places the nodes with higher correlation closer together (2.3 Network production and visualization).

### 3.2 Networks at low and intermediate correlation thresholds reveals processing-stage sample type modules

Next, we analyze correlation networks at low and intermediate correlation thresholds 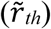, varying it from 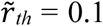 to 0.5, in steps of 0.1. The network at a given correlation threshold contains all the edges with correlation greater than or equal to the set threshold. Some of these networks are visualized in Figure 2. Panels A, B, C and D in Figure 2 show the network at correlation thresholds of 0.1, 0.3, 0.4 and 0.5, respectively. In panel A, the nodes belonging to Unfermented samples are seen little separated from the nodes belonging to the Fermented and Liquor samples. In panel B, the Unfermented samples are clearly separated from the Fermented and Liquor samples. Within the Fermented and Liquor samples little grouping starts to form. In panel C, the separation between the Fermented and Liquor samples becomes enough clear. And in panel D, all the three samples can be seen clearly separated from one another. This separation of samples first into two groups: (a) Unfermented, and (b) Fermented and Liquor samples, and then slowly into three groups: Unfermented, Fermented and Liquor samples, is in congruence with the earlier result seen in the structure of the correlation matrix heatmap shown in Figure 1A. Both Figure 1A and Figure 2B,C show that the liquor sample are more similar to the fermented samples than to the unfermented samples. This is in accordance with the fact that major chemical and physical changes in cocoa beans takes place during the processes of fermentation. A movie of the network as a function of progressively increasing the threshold is attached as supplementary information which clearly shows the evolving network and separation of samples belonging to different cocoa processing stage. Similar behavior is noted for the case of correlation network formed using the Pearson correlation coefficient (Supplementary Information file) however at different values of correlation threshold.

**Figure 2.**
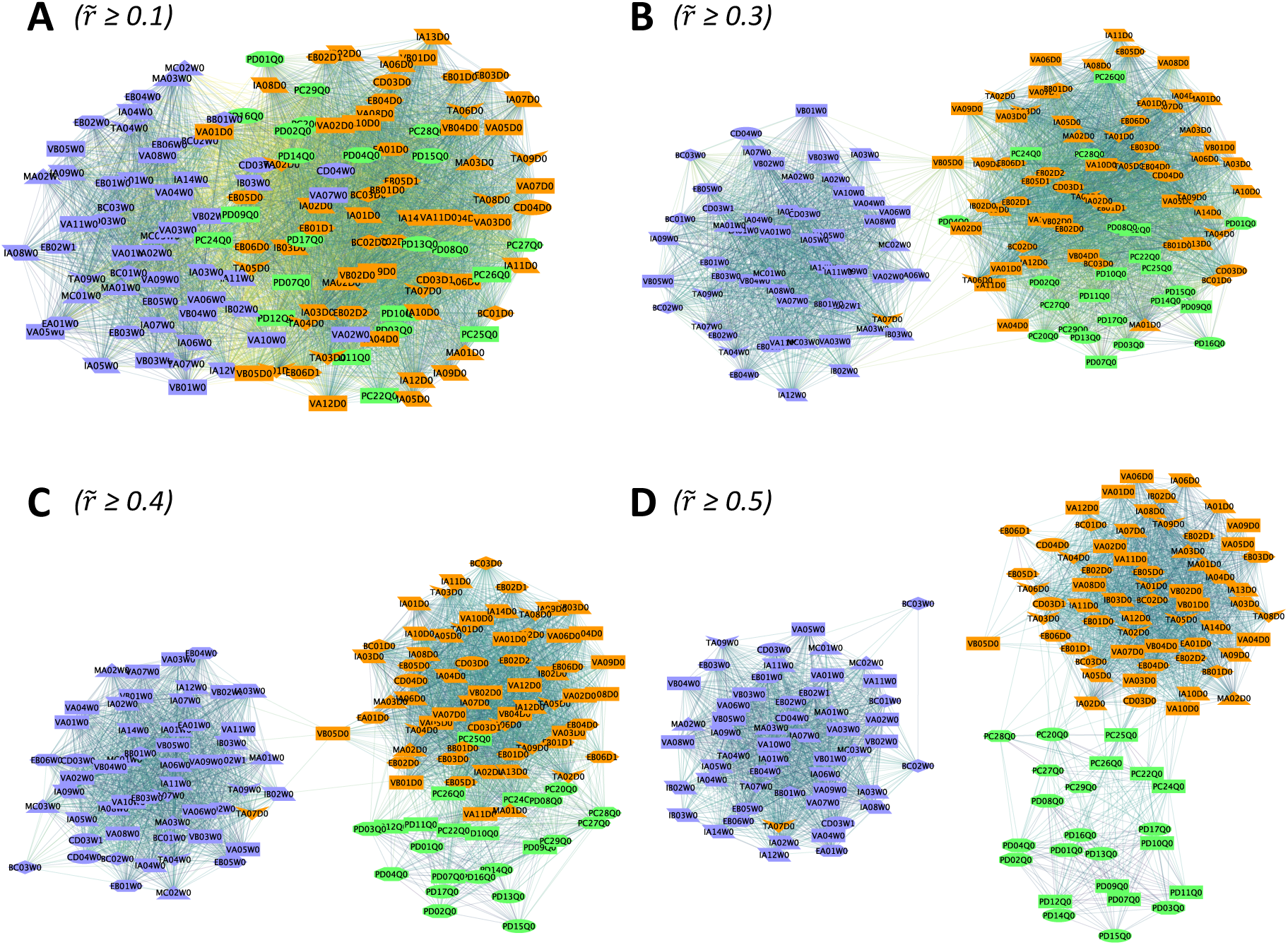
Processing-stage modules in low and intermediate threshold correlation networks. The figure reveals modules of samples belonging to the same cocoa processing-stage in a typical cocoa processing pipeline. **(A)** Network of LC-MS samples at a correlation threshold of 0.1 revealing separation of unfermented, fermented and liquor cluster. **(B, C)** Correlation thresholds 0.3 and 0.4. The separation between different processing-stage sample types improves. **(D)** Correlation threshold 0.5. Three groups of unfermented, fermented and liquor samples are clearly separated. The figure follows same legend as of Figure 1B**Error! Reference source not found.**. See supplementary information for a movie on evolving network as the correlation threshold is progressively increased.

### 3.4 Country enriched modules at high correlation thresholds

As the correlation threshold is further increased, the network breaks into various smaller connected components. The resulting individual connected components primarily have the processing-stage sample type. However, there are more than one component that belong to same color or sample type. This reveals the internal structure of the clusters of samples that initially grouped on the basis of their sample types. This additional sub-structure of the network reveals grouping which now is primarily governed by the samples belonging to same country of origin. This is shown in the networks in Figure 3 for correlation thresholds of 0.6, 0.7 and 0.8. Panels A and B provide a bird’s-eye view at respective thresholds, while panel C gives a detailed view. In contrast to the legend used in previous figures, we now color the nodes on the basis of countries for a quick comprehension of grouping on the basis of countries. The figure with the previous legend scheme is given as Supplementary Information. It can be seen from the figure that same color nodes tend to be present closer together. This feature is visible more in modules of smaller size, but it is also discernible in larger sized modules. We see that as the correlation threshold is further increased, most of the larger size modules break into smaller module, where nodes belonging to the same origin country are increasingly often connected. It should be noted that processing-stage and country of origin are only the major governing factors, on which grouping of samples is based. Other factors such as variety of cocoa hybrid, harvest season, geographical location and landscape of farm in the country etc, can begin to play an important role with increasing correlation threshold. Hence the clustering is not perfect. The other governing factors can potentially lead to finer sub-modular structures in the network. This situation is more likely to be evident at still higher correlation thresholds.

**Figure 3.**
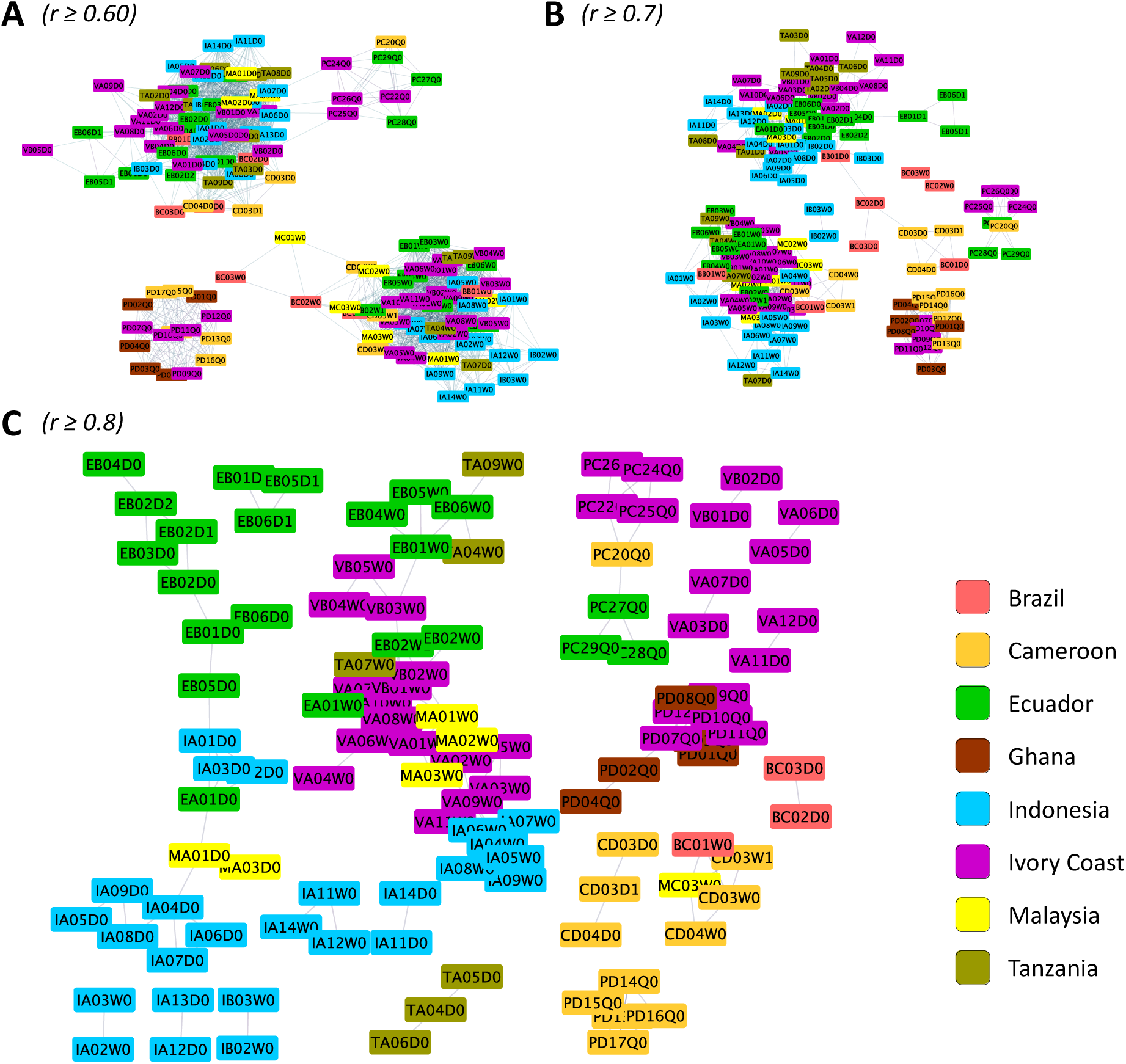
Country modules. The structure of correlation network of cocoa samples based on their LCMS profile at correlation thresholds of 0.80, 0.85 and 0.90. At these correlation thresholds, several modules with nodes belonging to the same country of origin are revealed. For a quick and better comprehension and unlike the legend of earlier correlation networks, in this figure different countries are represented through a different color. The networks with same thresholds but with previous annotation (i.e. of Figure 1 and Figure 2) is given in Supplementary Information for comparison. See supplementary information for a movie on evolving network as the correlation threshold is progressively increased.

As the correlation threshold is gradually increased, edges with correlation less than the threshold value are lost from the network. On one hand this leads to increased consideration of the edges with higher correlation in the determination of the layout of the network, while on the other this, naturally, leads to decrease in the number of edges, and when possible, also decreases the number of nodes in the resulting network, resulting in network breakage. The variation of number of edges and number of nodes connecting them is shown in the Supplementary Information file. In our networks here, only the edges greater than the set correlation threshold and corresponding nodes are present. A movie of the network as a function of progressively increasing the threshold is attached as supplementary information which clearly shows the evolving network and separation of samples belonging to different countries.

### 3.5 Similarity of nodes connected by an edge

As a node in our correlation networks has two attributes, namely the processing-stage sample-type and origin, we define two kinds of similarity for a pair of nodes connected by an edge: sample-type similarity and origin similarity. We define sample-type similarity as the fraction of edges in a network connecting nodes having the same processing-stage sample-type attribute, and origin similarity as the fraction of edges in a network connecting nodes which have same origin attribute. The sample-type and origin similarities as a function of correlation thresholds based on Spearman correlation networks are shown in Figure 4 (solid lines). They differ significantly from each other in terms of both the correlation threshold around which they start to rise and the manner in which they rise. The sample-type similarity starts to increase right from the smaller values of correlation thresholds itself and in a linear manner until it starts to saturate around a correlation threshold value of 0.5 to a similarity value close to 1. This is in agreement with the observed enhancement of the processing-stage sample type character of the network architecture right from the beginning of starting values of correlation threshold, to the almost full appearance of processing-stage sample type character at intermediate correlation threshold in large and small connected components (cf. Figure 2). The origin similarity remains almost constant and close to that of null model networks (orange dashed line) for a long range of correlation threshold (up to 0.5) suggesting a weak or almost negligible role in the clustering of nodes belonging to the same origin in the layout of network. Only when the correlation threshold is around 0.5, origin similarity starts to increase, suggesting this is the value of correlation threshold at which the contribution of origin effects start to contribute in clustering of nodes belonging to same origin begins. This clearly shows that the processing-stage sample type effect precedes the country effects, and the country effects are finer than the sample-type effect. The origin similarity increases exponentially and reaches a value close to 1. This implies that at higher threshold almost all edges connect nodes having same sample type and same country of origin.

**Figure 4.**
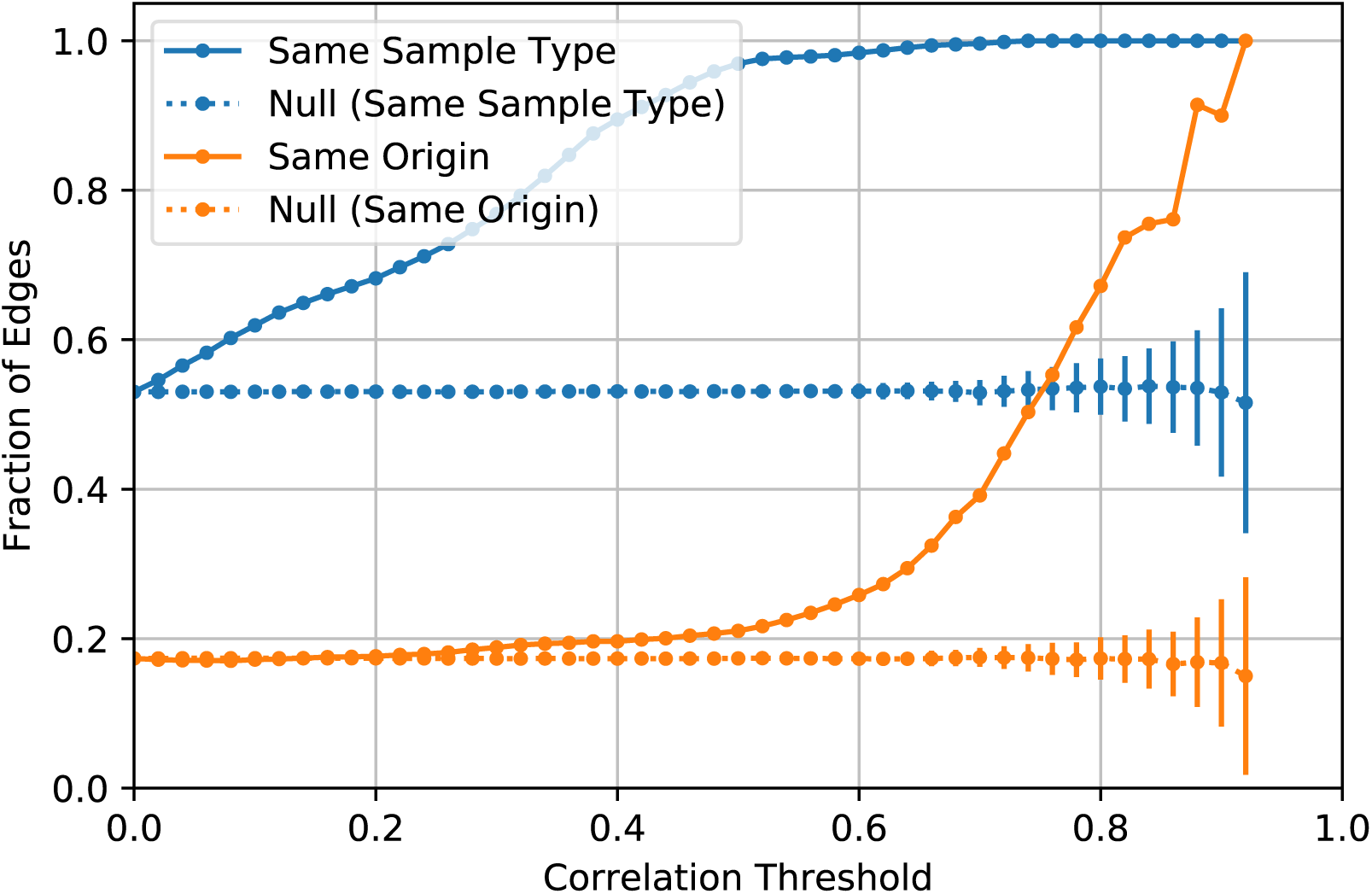
Connected nodes’ similarity. The sample-type similarity (blue line) starts to increase linearly right from smaller correlation threshold values, reaches close 1 around a correlation threshold value of 0.5. The origin similarity remains constant for a long range of correlation threshold (0, 0.50) and then increases exponentially. The dashed lines show corresponding similarities as expected from an ensemble of control networks.

The dashed lines along with error bars show similarity values and standard deviation expected from an ensemble of null model networks (control networks) obtained by randomizing edge weights in the original network (see 2.4 Null model network or control network). The difference between the similarity values from original network and that obtained null model networks points to the fact that the networks at higher correlation thresholds are enriched in edges that have high sample-type and origin similarity. The result corresponding to correlation network generated using Pearson correlation coefficient is given in Supplementary Information file. Both show similar behavior, although at slightly different correlation threshold value.

### 3.6 Closeness of thresholded networks to ideal networks

In this section, we quantify as a function of correlation threshold how accurately our networks represent the expected ideal networks of cocoa samples given their processing-stage sample types or country of origin. We consider two ideal networks, one each for the processing-stage sample type and country of origin. An ideal processing-stage sample type based network will have a link between a pair of its nodes only when both the nodes belong to the same processing-stage sample type, otherwise the link would be absent. Similarly, an ideal origin-based network will have a link between a pair of its nodes only when both the nodes belong to the same country of origin. Thus, in an ideal network based upon processing-stage sample type or country of origin a link is present only between nodes belonging to same sample type, or nodes belonging to same origin, otherwise there is no link between dissimilar nodes. After defining these ideal or true networks, we identify ‘true positive’ and ‘true negative’ links by comparing the links in the original network at a given correlation threshold (or thresholded network, for short) with the links in the ideal networks. A link is counted to be ‘true positive’ when the link is present both in the original network at the given threshold and the corresponding ideal network. A link is counted as ‘true negative’ when the link is absent both in the network at the given threshold and the corresponding ideal network. On the other hand, a link is defined as ‘false positive’ when it is present in the thresholded network but not in the corresponding true network, and ‘false negative’ when it is absent in the thresholded network but present the true network. An illustration of this scheme through a toy network is provided in Supplementary Information file. Using these terms, we define accuracy α as the fraction of ‘true positive’ and ‘true negative’ links in an original thresholded network. Accuracy quantifies how close a thresholded network is to the ideal expected network.

We find that with increasing correlation thresholds the network becomes closer to the expected true network as demonstrated by increasing values of accuracy for both processing-stage sample type and country of origin Figure 5 (Spearman correlation network; Pearson correlation case in Supplementary Information file). Further, in the region of low correlation threshold the character of the network is closer to that of the expected true network for the processing-stage sample type attribute, and in the region of higher correlation threshold the character of the network is closer to that of the expected true network for country of origin attribute. This result is in agreement with the previous results with formation of processing-stage sample type clusters at lower and intermediate correlation thresholds and of country-based clusters at high correlation thresholds.

**Figure 5.**
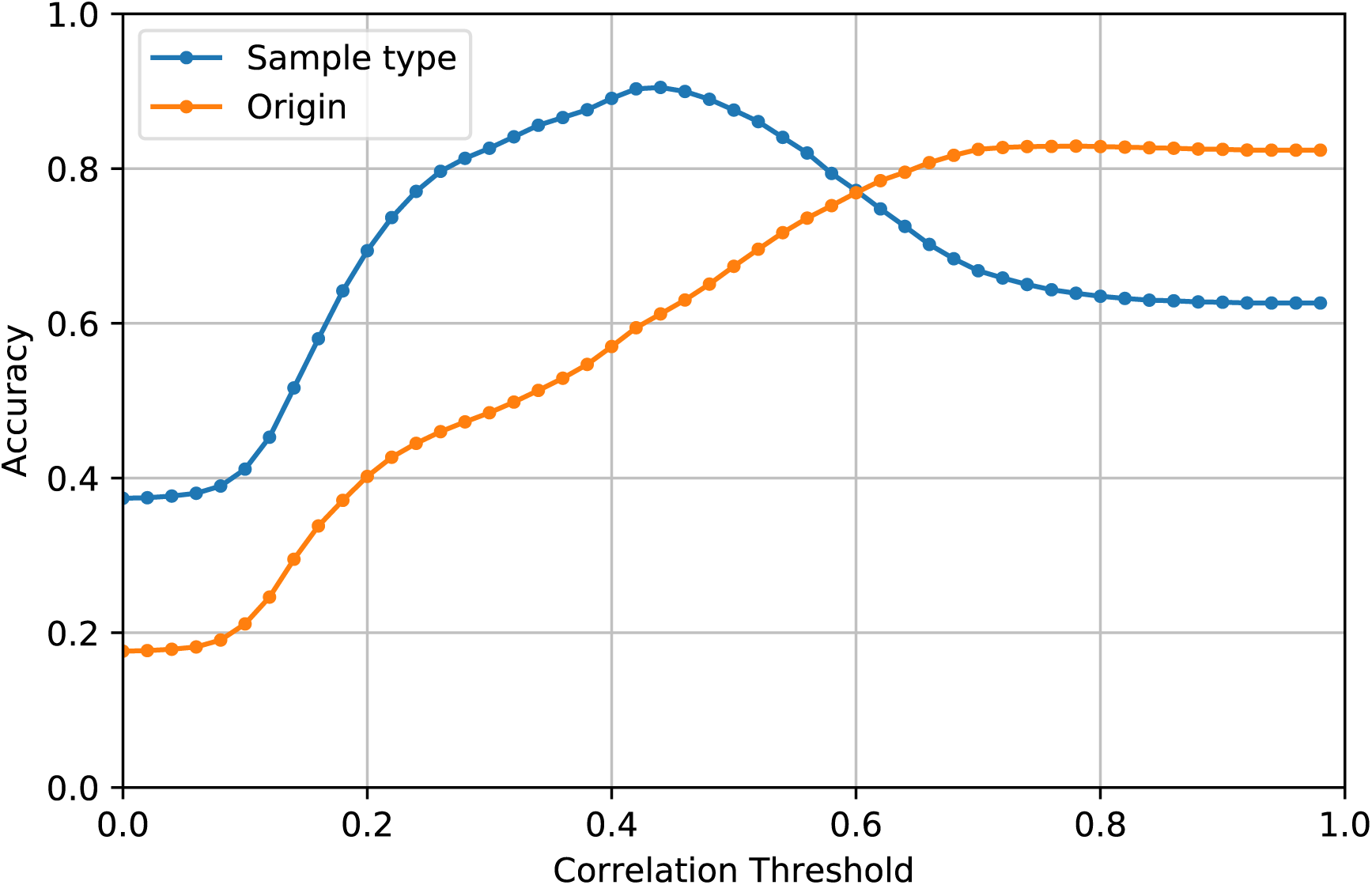
Accuracy of links in thresholded correlation networks, or closeness of a threholded correlation network to expected ideal network based on sample type or origin attributes of cocoa samples. As the correlation threshold increases the threshold networks become closer to their ideal counterparts. In regions of lower correlation threshold, the thresholded networks are describe more the sample type character of the network than the origin type character. In regions of higher correlation threshold, opposite is true and the thresholded networks are closer in their character to the origin attribute of LC-MS samples. This is coherent with the network pictures at various threshold seen in earlier figures.

## 4. Conclusions and Discussion

We have introduced a new approach for studying grouping in cocoa samples using their LC-MS profile. This new approach is often called ‘network science’, and it already benefits a multitude of scientific disciplines. Few cases also exist where network approach has been successfully applied in food science for different purposes (Hochberg et al., 2013; Ursem et al., 2008; Wang et al., 2017), however, to the best of our knowledge, we apply it for the first time to study the classification of cocoa samples based upon their LC-MS profiles.

Classification of cocoa samples on the basis of their country of origin has been found challenging with limited success obtained in cases with the number of countries being few or the origin being on continental scale. Differences in unfermented and fermented samples can be easily seen by simply finding the Spearman correlation between the cocoa samples using their LC-MS profiles (cf. Figure 1). The liquor samples are closer to the fermented samples. However, differentiation on the level of country of origin is only revealed upon further analysis. We make a correlation network using the correlation matrix for cocoa samples, and show that systematic variation of a single parameter, namely correlation threshold, can be used to reveal grouping of cocoa samples on the basis of processing-stage, viz. unfermented, fermented and liquor, and country of origin. In the low and intermediate ranges of correlation threshold processing-stage sample type clusters are revealed, and in the higher range of correlation threshold the clustering of cocoa samples on the basis of country of origin is witnessed. We present our results both qualitatively (cf. Figure 2 and Figure 3) and quantitatively (cf. Figure 4 and Figure 5). Besides a successful working approach, our work shows that differentiation of cocoa samples on the level of country of origin is on a more subtle level than their differentiation on the basis of processing-stage sample types.

It is worth comparing our approach to an often-used method in similar situation—the principal component analysis (PCA). PCA projects the samples into a lower dimensional space whose axes represent highest possible variation on the basis of the features in the dataset used in the analysis. Often it turns out that this analysis is also able to provide us a view in which samples with different classes well separated. However, there is no binding reason for it to be so, as PCA focuses on maximizing variation amongst the samples on the basis of their features and not clustering them per se. Further, only truncated amount of information can be used to visualize the samples as we are limited to a maximum of three dimensions. On the other hand, in a correlation network information from all features (compounds used to calculate correlation) is present. Further, one is able to look at the structure of the network at the level of different amount of information by pruning the network thereby keeping low/high correlations as per need. In this sense, the approach of correlation networks is more sophisticated than that of PCA, omitting PCAs basic philosophy of data reduction.

Our study takes into consideration two factors on which cocoa samples may primarily differ: processing-stage and country of origin. However, it is worth noting these are not the only governing factors that affects similarity of cocoa samples. Many other factors such as variety of cocoa hybrid, soil, climate, terrain, harvesting season, farming practices etc. also have significant effects (Acierno et al., 2016; Adeniyi et al., 2019; Arévalo-Hernández et al., 2019; Ehiakpor et al., 2016; Kongor et al., 2016). It would be interesting to consider some of these factors in future works and see in what range of correlation threshold these effects start to matter, or can the inclusion of these additional factors give more clear modules of cocoa samples. Besides providing a new approach to study similarity in cocoa samples, our approach can be a compliment to the traditional approaches in this field.

## Supporting information

video_spearmanCorrelation_sampleType

## Acknowledgements

We thank the whole COMETA team at Jacobs University which generated and gathered the LC-MS data over a period of five years, in particular Nina Böttcher and Britta Brehends for their excellent support in sample procurement, preparation and running experiments. SK is thankful to Johannes Falk and Piotr Nyczka for various stimulating and insightful discussions. This work was funded by the COMETA Project, which is financially supported by Barry Callebaut AG.

