## Supporting Information for "Correlation network analysis based on untargeted LC-MS profiles of cocoa reveals processing stage and origin country"

#### 1 Pearson correlation

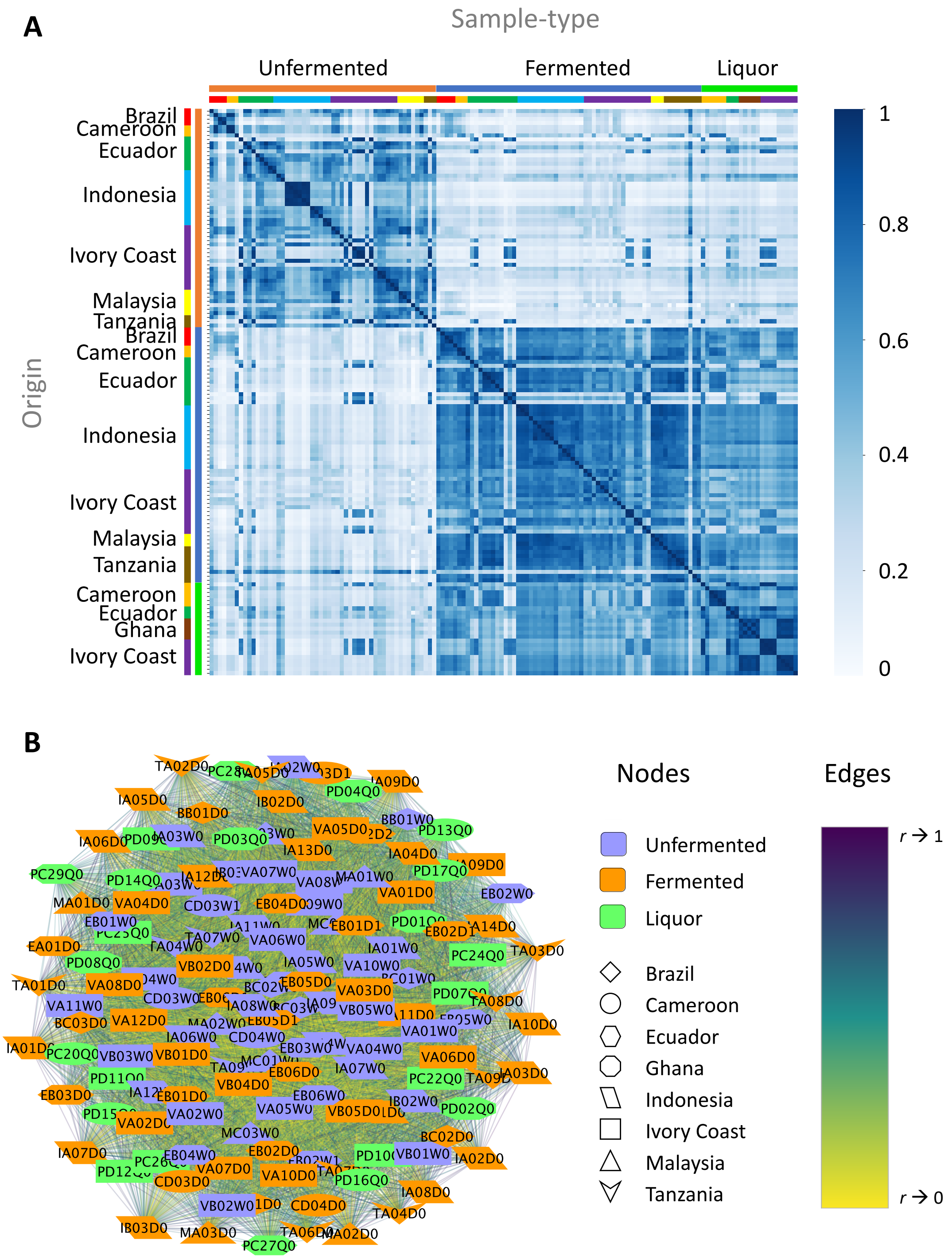

**Figure 1 Correlation between cocoa samples.** Same as Fig. 1 in main text, but using Pearson correlation instead of Spearman correlation. **(A)** Correlation heatmap**.** The distinction between the Fermented and Liquor samples is lesser compared to that brought out using Spearman correlation. Unfermented samples are easily distinguishable from the Fermented and Liquor

samples. (**B)** Correlation Network. The correlations are computed using Person correlation coefficient.

#### 2 Networks at low and intermediate correlation using Pearson correlation

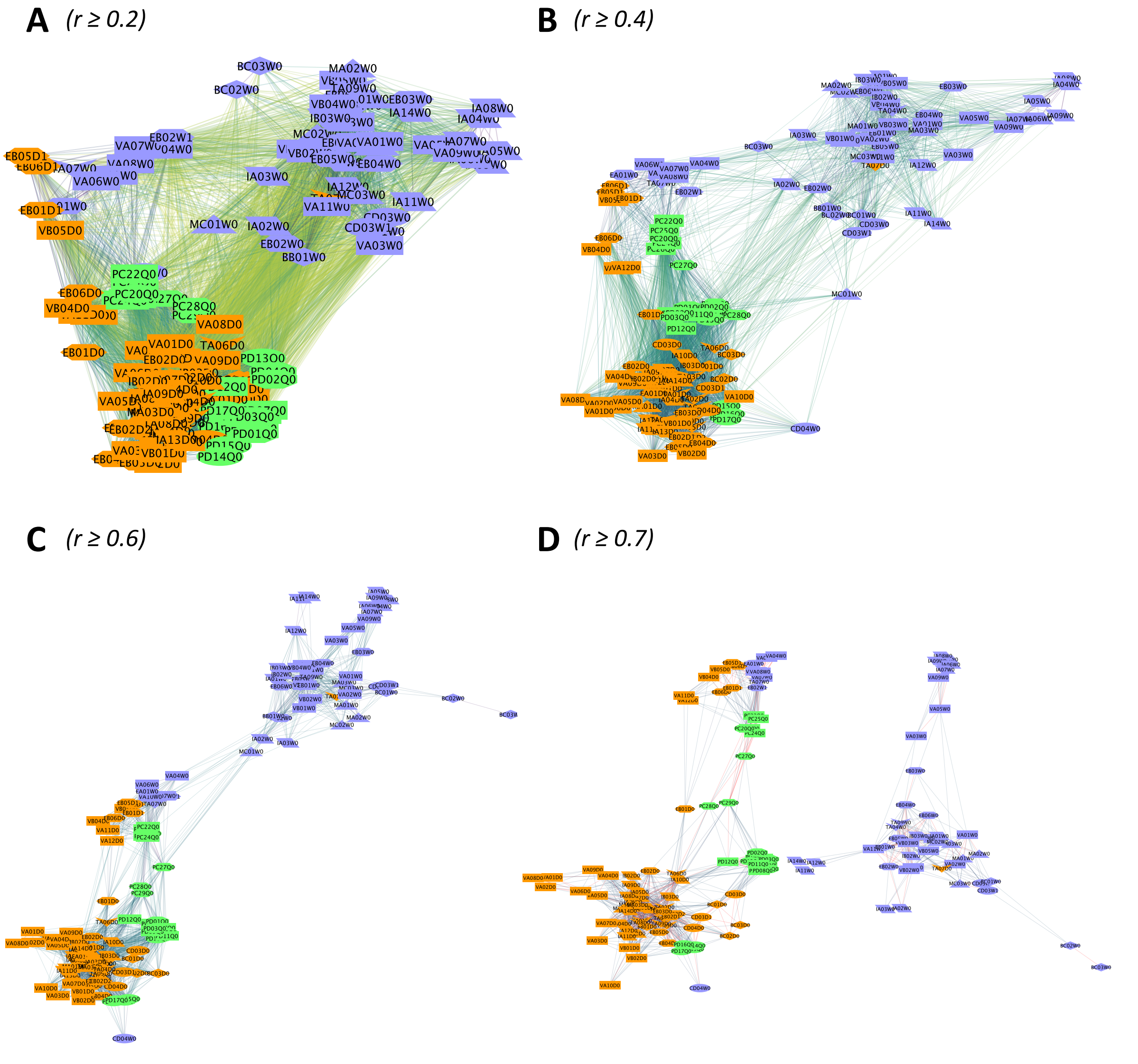

**Figure 2 Correlation at different correlation thresholds.** Similar to Fig. 2 in main text, but the Pearson correlations are used to make the network. The revelation of substructure of the network showing Unfermented, Fermented and Liquor occurs at different threshold. Nevertheless, the substructures are clearly revealed.

### 3 Country enriched modules at high correlation threshold

#### 3.1 Using Spearman correlation

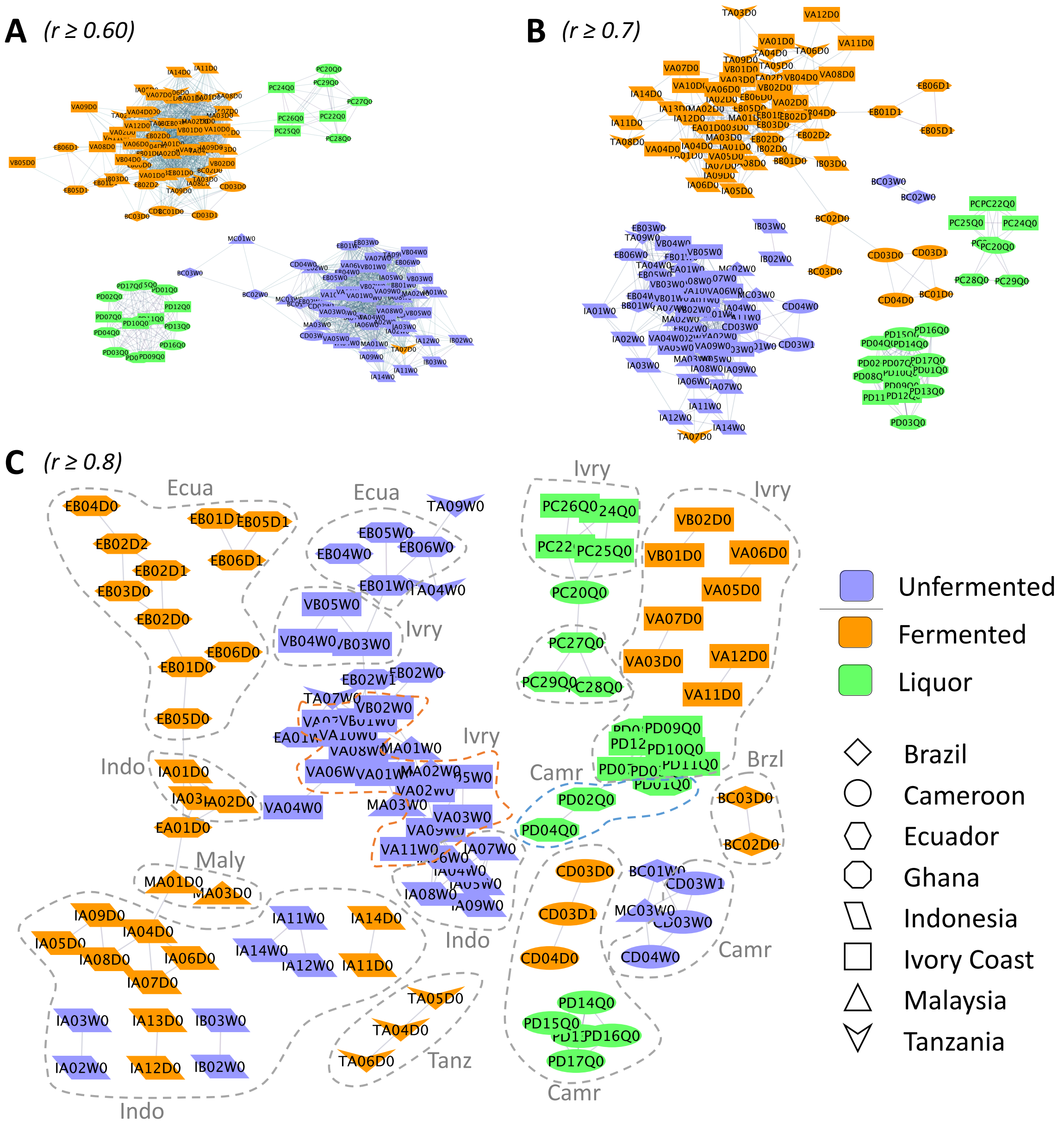

**Figure 3 Country modules revelated at higher correlations:** Same as Fig. 3 in main text, but with node color representing sample-type and node shape representing countries of origin of cocoa sample. For easy comprehension of revealed grouping on the basis of origin, nodes belonging to the same country have been demarcated with dotted lines, and labelled with country legend—*Brzl*: Brazil, *Camr*: Cameroon, *Ecua*: Ecuador, *Ghna*: Ghana, *Indo*: Indonesia, *Ivry*: Ivory Coast, *Maly*: Malaysia; *Tanz*: Tanzania. The color and shape of the dotted lines are only

for the purpose of clear comprehension; it does not have any further connotation. The electronic version of the image can be zoomed in for details.

#### 3.2 Using Pearson correlation

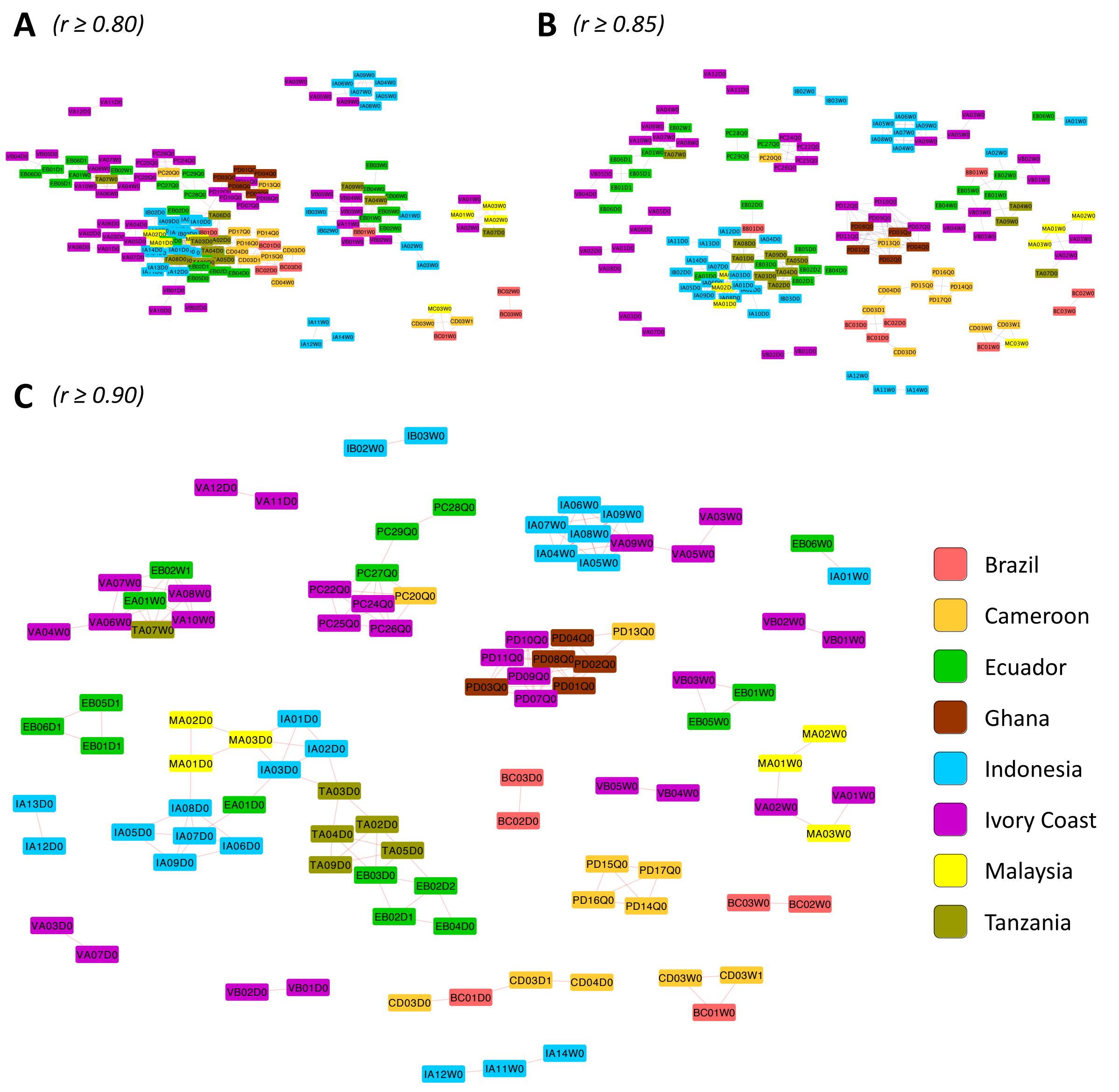

**Figure 4 Country modules revelated at higher correlations**: Same as Fig. 3 in main text, but with Pearson correlation instead of Spearman correlation.

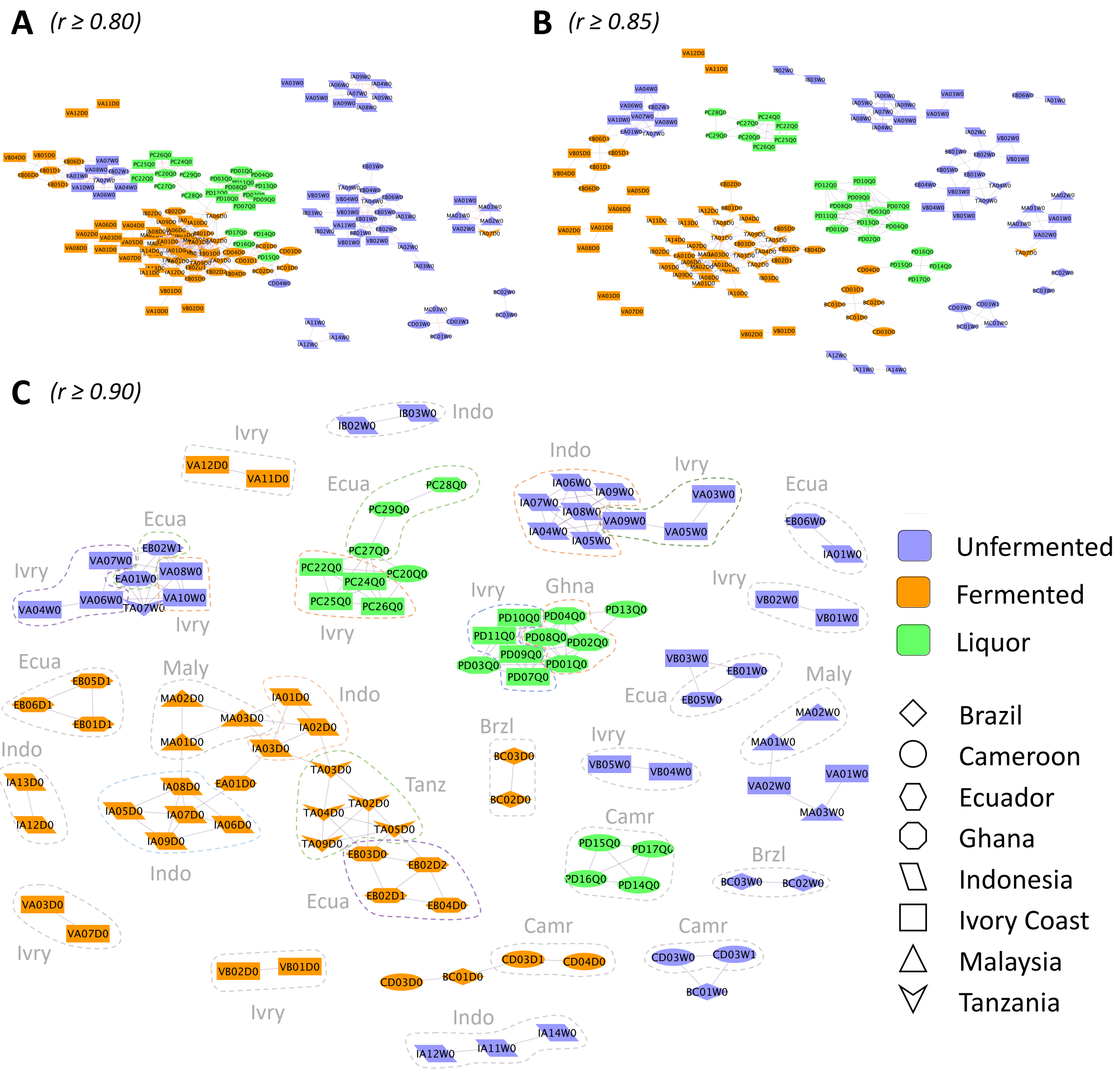

**Figure 5 Country modules** Same as Fig. 4 in the supplementary text i.e., using Pearson correlation coefficient, and with node color representing sample-type and node shape representing countries of origin of cocoa sample. For easy comprehension of revealed grouping on the basis of origin, nodes belonging to the same country have been demarcated with dotted lines, and labelled with country legend—*Brzl*: Brazil, *Camr*: Cameroon, *Ecua*: Ecuador, *Ghna*: Ghana, *Indo*: Indonesia, *Ivry*: Ivory Coast, *Maly*: Malaysia; *Tanz*: Tanzania. The color and shape of the dotted lines are only for the purpose of clear comprehension; it does not have any further connotation. The electronic version of the image can be zoomed in for details.

##### 3.3 Number of nodes and edges as a function of correlation threshold in networks using Spearman and Pearson correlation

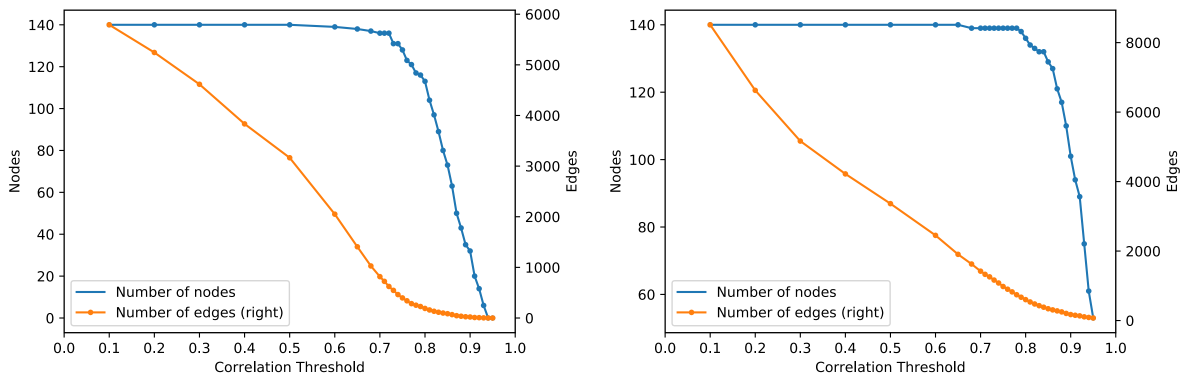

**Figure 6 Number of Edges and nodes at different correlation thresholds in correlation network made using Spearman correlations. (Left panel)** Nodes and edges in network made using Spearman correlations. As the correlation threshold in increased, number of edges in the resulting networks drops sharply. The number of nodes remains almost constant for a wide range of the correlation threshold (0, 0.6), changes slowly in range (0.6, 0.75), and afterwards witness a sharp fall. **(Right panel)** Nodes and edges in network made using Pearson correlations. The number of nodes remains almost constant for a wide range of the correlation threshold (0, 0.7), changes slowly in range (0.7, 0.8), and afterwards witness a sharp fall.

### 4 Similarity of nodes connected by edges in networks made using Pearson correlation

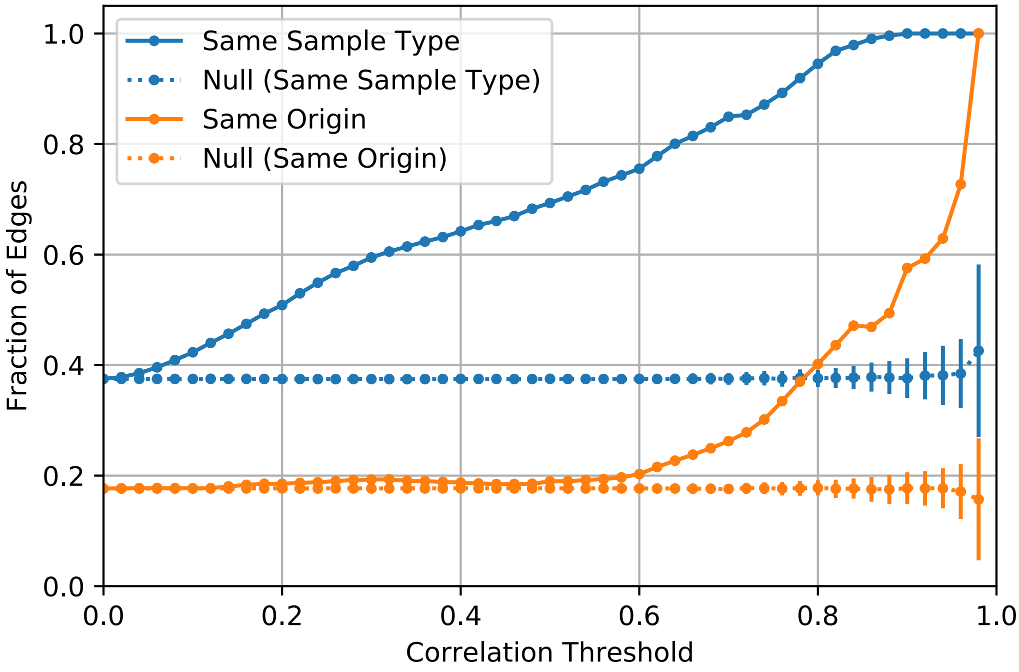

**Figure 7 Connected nodes' similarity in network made using Pearson correlation coefficient.** Same as Fig. 4 in the main text, but in the correlation network made using the Pearson correlation instead of Spearman correlation. Qualitatively, the behavior is same in both cases, i.e., Spearman correlation network and Pearson correlation network. In the latter case, i.e., Pearson correlation network, same similarity values are reached at higher correlation thresholds. In both cases, sample-type similarity value remains high, and origin similarity catches up at high correlation thresholds.

### 5 Accuracy of links in thresholded correlation networks

#### 5.1 Toy network illustrating accuracy concept

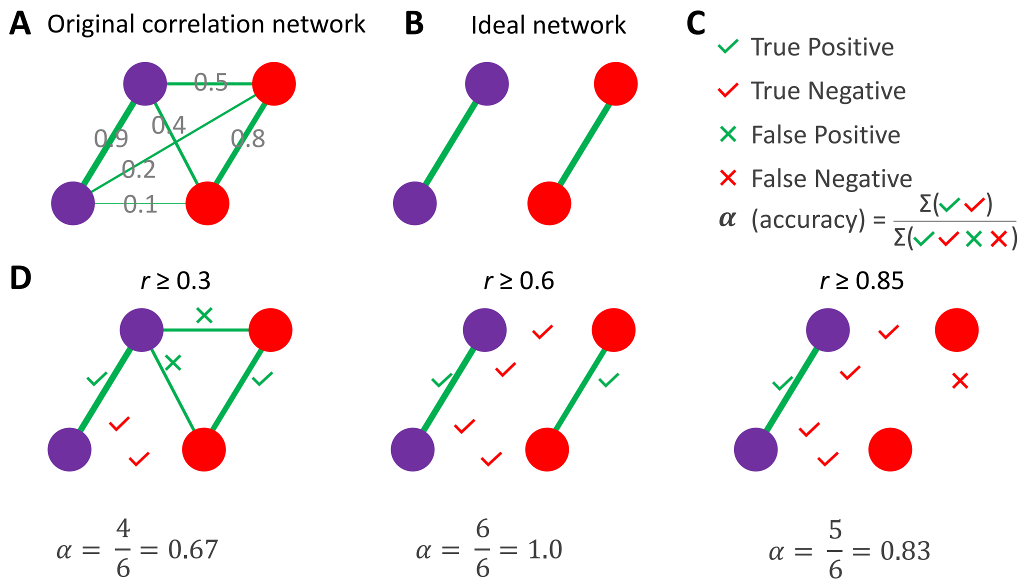

**Figure 8 Toy example illustrating calculation of accuracy of links using original and expected ideal network at different correlation thresholds.** Same color of the node may be taken as representing one of the either attribute of cocoa samples: sample type or country of origin. **(A)** The original correlation network obtained after finding correlation between cocoa samples. The correlations between nodes are mentioned on respective links/edges. **(B)** The expected ideal network: it has links only between same color nodes (same attribute: sample-type or same origin). **(C)** Legend for the link type in thresholded networks. True positive: link present in both thresholded and ideal network; true negative: link absent in both thresholded and ideal network; false positive: link present in thresholded network but absent in ideal network; and false negative: link absent in thresholded network but present in ideal network. Accuracy: fraction of true positive and true negative links in the thresholded network. **(D)** Example networks at different [correlation] thresholds and accuracy of the links, or alternatively, closeness of the original thresholded network to the expected ideal network.

#### 5.2 Accuracy of link in correlation network made using Pearson correlation as a function of correlation thresholds

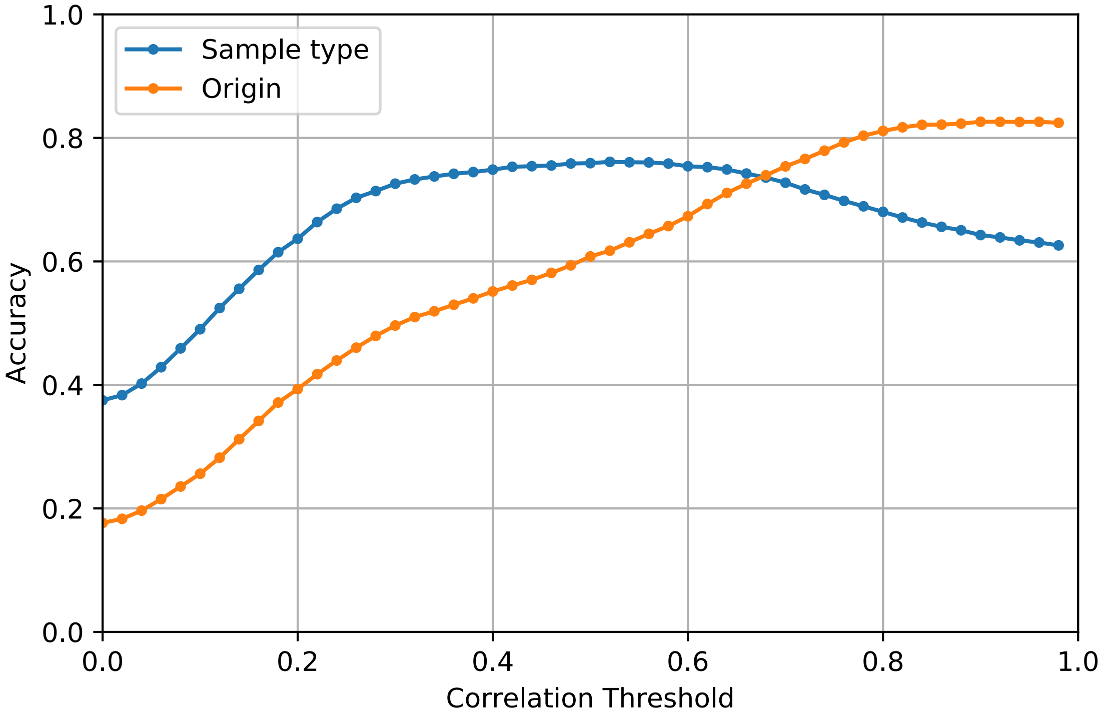

**Figure 9 Accuracy of links in thresholded correlation networks, or closeness of a threholded correlation network to expected ideal network based on sample type or origin attributes of cocoa samples.** Same as in Fig. 6 in main text, but in the correlation network made using Pearson correlation instead of Spearman correlations. Compared to the case of Spearman correlation network in Fig. 6 in main text, here the plateau for sample-type remains lower than 0.8. However, in both cases, the result qualitatively remains the same: at lower threshold the correlation networks are closer in character to the sample type feature of samples, while at higher thresholds the correlation networks are closer in character to the origin feature of samples.
